# Spatiotemporal distribution and species diversity of pathogenic *Vibrios* in estuarine recreational waters of southeast Louisiana

**DOI:** 10.1101/2025.07.16.665201

**Authors:** Annika Nelson, Fernanda Mac-Allister Cedraz, Katie Vigil, Joshua Alarcon, Tiong Gim Aw

**Affiliations:** Department of Environmental Health Sciences, Celia Scott Weatherhead School of Public Health and Tropical Medicine, Tulane University, New Orleans, Louisiana, USA

**Author notes:** Corresponding author: Tiong Gim Aw Mailing address: Department of Environmental Health Sciences School of Public Health and Tropical Medicine Tulane University, 1440 Canal Street, Suite 2100 New Orleans, LA 70112, USA.

**Keywords:** *Vibrio cholerae*, *Vibrio vulnificus*, *Vibrio parahaemolyticus*, recreational water, environmental reservoir, nanopore sequencing

## Abstract

*Vibrio* bacteria occur naturally in brackish water and can cause illnesses through recreational water exposure. *Vibrio* infections have shown a notable increase in recent years worldwide. In this study, bacterial culture and molecular methods were used to assess the prevalence and diversity of *Vibrio* species in the estuary of Lake Pontchartrain, the second largest inland brackish body in the United States. Water samples (n= 101) were collected from 9 recreational sites from November 2023 to November 2024. During the summer months (June, July, and August), the average *Vibrio* species. concentration was 5.2 × 10^4^ colony-forming unit (CFU)/L. While in the winter months (December, January, and February), the average *Vibrio* spp. concentration was 3.2 × 10^3^ CFU/L. Likewise, the temperature differed between summer and winter, with the average water temperatures being 30.39 °C and 14.45 °C, respectively. Linear modelling showed water temperature and salinity were found to be significant (p < 0.05) predictors of *Vibrio* concentrations from both culture methods and quantitative PCR, while precipitations were only significant for bacterial culture. The *toxR* genes of *Vibrio cholerae*, *Vibrio vulnificus*, and *Vibrio parahaemolyticus* persisted throughout the year, and 14.8% (n=9) of sequenced samples were identified as the O139 serotype of *V. cholerae.* Bacterial isolate sequencing revealed more than 100 *Vibrio* species in the lake with *V. cholerae, V. vulnificus,* and *V. mimicus* making up the largest proportion of the community. Continuous environmental monitoring of *Vibrio* is warranted in informing public health preparedness and expanding our understanding of the ecology of this pathogen.

**Importance:** Globally, the diverse bacterial genus *Vibrio* is an important group of pathogens in coastal water environments. These bacteria are responsible for waterborne and seafood-borne illnesses as well as skin infections from recreational activities. Despite the rising incidence of *Vibrio* infections, routine monitoring of *Vibrio* species (spp.) in the environment remains limited. This gap hinders our understanding of their distribution, especially in estuarine areas, and potential public health risks linked to recreational activities. This study employed multiple techniques including bacterial culture, quantitative PCR, and genome sequencing to assess the prevalence, distribution, and diversity of *Vibrios* in the estuary. This study shows that *Vibrio cholerae* were prevalent in recreational water of a non-endemic region. The findings underscore the need for regular monitoring of *Vibrio* levels in recreational water and educating the public on the risks.

## 1 Introduction

Many species (spp.) of *Vibrio* in coastal environments can cause serious human infections, representing a growing global health concern linked to increasing seawater temperatures (Deeb et al., 2018). The *Vibrio* genus includes over 100 species of gram-negative bacteria that are found in a wide range of aquatic environments, from freshwater to marine water (Brumfield, Chen, et al., 2023) . *Vibrio* spp. are naturally present in the environment and are known to grow best in slightly salty waters, ranging from 1 to 35 parts per thousand (ppt), with temperatures between 25 to 30 °C (Martinez-Urtaza et al., 2008; Sheikh et al., 2022).

Studies have shown that environmental factors such as water temperature, salinity, and nutrient load can affect the type and number of *Vibrios* that are present in an aquatic system (Brumfield, Chen, et al., 2023; Froelich et al., 2019; Leal Filho et al., 2022). The three species of concern have different salinity ranges for maximum growth, with *V. cholerae* grows optimally in water with low salinity, ranging from 2 to 8 ppt. The optimum range of salinity for the growth of *V. vulnificus* is from 5 to 25 ppt, and *V. parahaemolyticus* is near 25 ppt (Louis et al., 2003; Martinez-Urtaza et al., 2008; Randa et al., 2004). In the Chesapeake Bay, total dissolved solid (TDS) values between 10 to 15 g/L were associated with the increase of *Vibrio* detection (Brumfield, Chen, et al., 2023). Additionally, *Vibrio* spp. can enter a viable but non-culturable (VBNC) state in response to stressful conditions such as temperature fluctuations and other environmental factors (Balagurusamy et al., 2024; Colwell & Huq, 1994). Unlike dead cells, VBNC cells maintain low but detectable metabolic activity (Balagurusamy et al., 2024). This state enables *Vibrio* spp. to persist during colder months, and when conditions become favorable, they can resuscitate, regaining normal metabolic function, culturability, and virulence (Balagurusamy et al., 2024).

Of the many species of *Vibrio* that inhabit water systems around the world, 12 are pathogenic to humans. *Vibrio* spp. are most commonly associated with human gastrointestinal illness including *V. cholerae*, *V. vulnificus*, *V. parahaemolyticus,* and *V. alginolyticus* (Jones, 2014; Robert-Pillot et al., 2014). *V. cholerae*, specifically in the O1 or O139 serotypes produce a toxin that causes secretion of water and electrolytes from cells, resulting in potentially-deadly, watery diarrhea (Vanden Broeck et al., 2007). Cases of non-cholera *Vibrio* spp. (for example, *V. parahaemolyticus, V. alginolyticus* and *V. vulnificus*) infections are often classified as vibriosis, but different species can cause different symptoms or severity of infection (Baker-Austin et al., 2018). In humans, these bacteria can cause primary sepsis, wound infection including skin necrosis, and gastrointestinal illness (Chuang et al., 1989). *V. vulnificus*, the leading cause of seafood-associated fatality in the United States (U.S.), has an 80% rate of hospitalization and a case-fatality ratio greater than 30% (Heng et al., 2017; Newton et al., 2012). From the period of 1996 to 2010, *V. parahaemolyticus* was the most commonly reported *Vibrio* spp. infection in the U.S., but it had a case-fatality ratio less than 1% (Newton et al., 2012).

The Centers for Disease Control and Prevention (CDC) estimates that there are 80,000 cases of vibriosis in the U.S. each year, but many of these go unreported or misdiagnosed (CDC, 2024). In 2019, there were 2,685 confirmed, non-cholera vibriosis cases and 11 cholera cases in the U.S. Of these, there were 159 cases caused by *V. vulnificus*, with 52% of those cases being attributed to Gulf Coast states, including 12 cases in Louisiana (CDC, 2019). The Louisiana Department of Health reported in the 20-year period from 1988 to 2018, there were 628 reported vibriosis cases, and 35.8% of cases were attributed to skin exposure from either saltwater or freshwater sources (*Vibrios Annual Report*, 2018).

Lake Pontchartrain is an estuarine embayment that borders the northern side of the city of New Orleans and opens into the Gulf of Mexico via Lake Borgne to the east. It is a popular site for swimming, boating, fishing, and other recreational activities. Previous studies in the Gulf of Mexico showed that *Vibrios* were present in surface waters with fluctuating concentrations based on environmental conditions (Bernáldez-Sarabia et al., 2021; Johnson et al., 2010; Nigro et al., 2011). While these studies have looked at *Vibrio* spp. communities in the Gulf of Mexico, none of them have used a combination of methods, including bacterial culture, quantitative polymerase chain reaction (qPCR), and long-read sequencing methods to characterize the *Vibrio* communities including both culturable and VBNC bacteria. Furthermore, as intense storms and climate fluctuations affect water systems around the world, it is important to monitor the changing behavior of important pathogens like *Vibrio* spp. (Leal Filho et al., 2022). Therefore, the objectives of this study were to (i) use bacterial culture methods and qPCR to identify the presence of *Vibrio* spp. of public health concern in a recreational estuary; (ii) determine the effect of environmental variables (temperature, salinity, dissolved oxygen, fecal indicator bacteria) on spatiotemporal distribution of *Vibrio* spp.; and (iii) use a long-read nanopore sequencing method to assess the diversity of viable *Vibrio* spp. and serotypes in surface water.

## 2 Materials and Methods

### 2.1 Site description

Lake Pontchartrain is a brackish estuarine embayment located in southeastern of Louisiana (Figure 1). It is the second largest inland estuary in the U.S., covering an area of 630 square miles (1,631 square km). The lake is an important resource, supporting agricultural and aquacultural activities, fishing tourism and recreation (EPA, 2025). Water samples were collected at nine recreational sites from November 2023 to November 2024. Over the course of the study, 101 surface water samples were collected.

**Figure 1.**
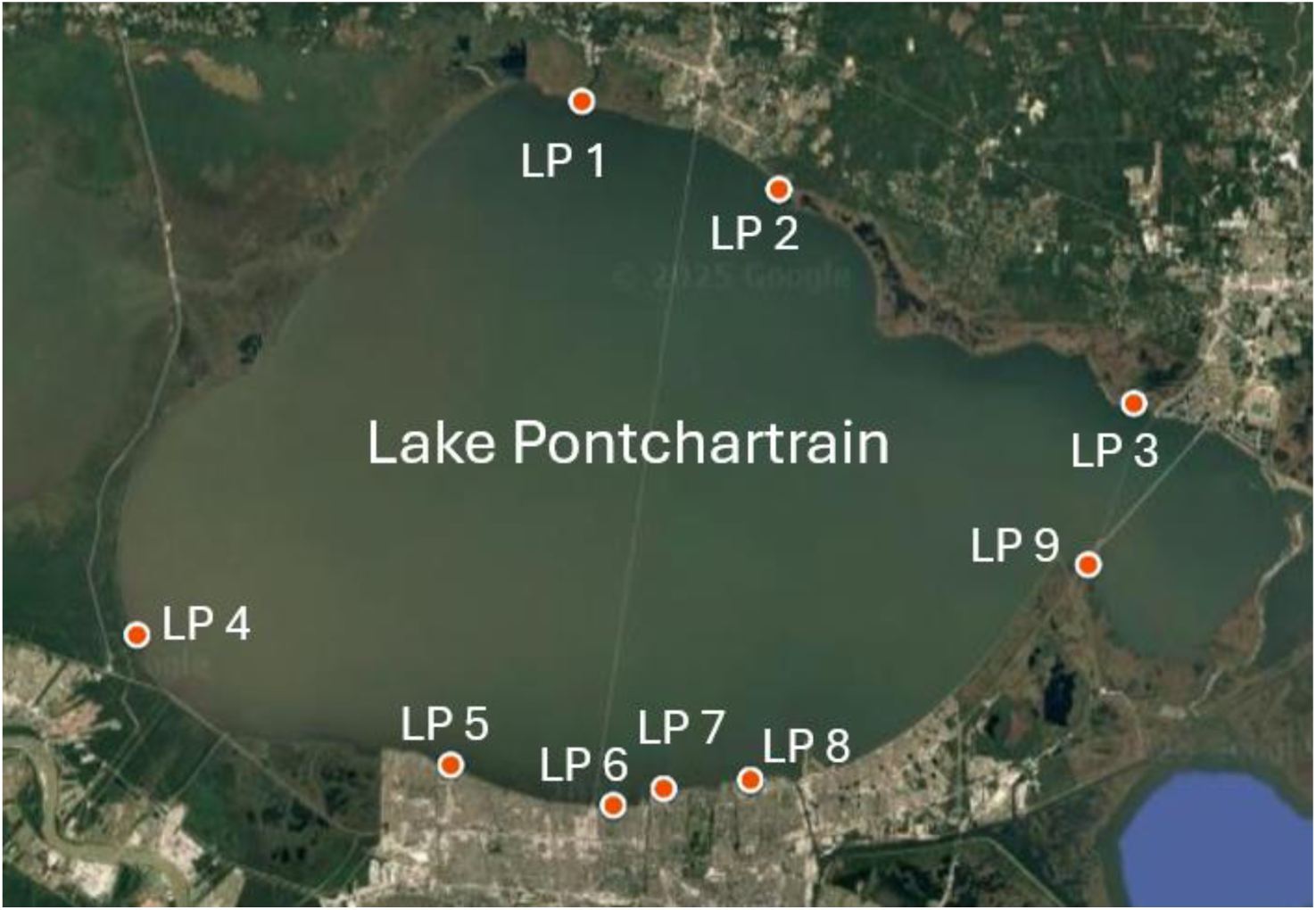
Map showing sampling sites along the shore of Lake Pontchartrain (LP). The LP 1 - LP 3 are sites located on the north shore of the lake, and LP 4 – LP 9 are the south shore sites.

### 2.2 Sample collection and water quality parameter measurements

During the study period, each site was sampled once a month. Two sampling trips were conducted each month, each trip collecting samples from the north shore of the lake (LP 1- LP 3) and south shore of the lake (LP 4- LP 9). Exceptions included December 2023 when no samples were collected, March 2024 when south shore samples were not collected, and November 2024 when LP 1 was not collected due to flooding. At the time of sample collection, a multiparameter weatherproof meter (HANNA Instruments, Smithfield, RI) was used to measure pH, dissolved oxygen (DO), salinity, and total dissolved solids (TDS). Precipitation data was obtained from the United States Geological Survey (USGS) National Water Information System at station #302415090091500 in Madisonville, Louisiana, U.S. A one-liter grab sample of water was collected from each site using a sterile bottle (Thermo Fisher Scientific, Waltham, MA). Water samples were stored in a cooler with ice packs during transport to the lab, where they were held at 4 °C until processing (<48 hours). Upon return to the lab, enzyme-based methods were used to assess the presence of fecal indicator bacteria. Briefly, the Colilert-18 reagent (IDEXX Laboratories, Westbrook, ME) was added to 100 mL of sample in a sterile sample vessel and mixed well to dissolve. The mixture was poured into an IDEXX Quanti-tray 2000 and incubated for 18 – 22 hours at 35 °C (±0.5 °C). Following incubation, wells showing yellow coloration (total coliforms) and fluorescence under 365nm UV light (*Escherichia coli*) were counted and interpreted using the IDEXX Most Probable Number (MPN) table, with results reported as MPN/100mL in accordance with manufacturer’s instructions. Similarly, for enumerating enterococci, 100 mL of sample was mixed with the Enterolert reagent in a sterile sample vessel (IDEXX Laboratories, Westbrook, ME), and incubated for 24-28 hours at 41 °C (±0.5 °C). After incubation, positive wells for enterococci (fluorescent under 365nm UV light) were counted and interpreted using the IDEXX Most Probable Number (MPN) table, with results reported as MPN/100mL in accordance with manufacturer’s instructions.

### 2.3 Enumeration of *Vibrio* spp. using culture-based assay

Within 48 hours of collection, water samples were filtered for use in a bacterial culture-based assay. Water samples were diluted with 1X sterile phosphate-buffered saline (PBS) to get the sample plates within the countable range. For each sample, a 1:10 (vol/vol) and 1:50 (vol/vol) 1X PBS dilution was made to get a total volume of 100 mL for filtration. Both dilutions and 100 mL of undiluted sample were filtered in duplicate on 0.47 mm diameter mixed cellulose ester (MCE) gridded filters with 0.45 μm pore size (Pall Corporation, Port Washington, NY). For each filtration, a negative control using 100 mL of sterile 1X PBS was included. Filters were placed on Thiosulfate-Citrate-Bile Salts-Sucrose (TCBS; Cat. No. 265020, BD Difco™) agar and incubated at 37 °C for 18-24 hours. After incubation, the colonies were counted. The countable range was <200 colonies per plate and the duplicate plates were averaged. The concentration of *Vibrio* spp. in the sample measured in colony forming units (CFU) per liter (CFU/L) was calculated by multiplying the number of colonies times the dilution factor and dividing by the volume filtered.

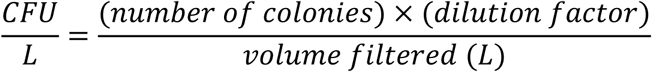

### 2.4 Confirmation of *Vibrio* species using Polymerase Chain Reaction

The TCBS agar is a highly selective medium for the isolation of different *Vibrio* spp., inhibiting gram-positive bacteria via bile salts and supporting *Vibrio* growth under alkaline conditions. It differentiates species by sucrose fermentation and hydrogen sulfide production, resulting in distinct colony colors and morphologies. *V. cholerae* formed large yellow colonies *V. vulnificus* formed yellow-green colonies, whereas *V. parahaemolyticus* produced blue to green colonies.

Therefore, yellow and green colonies from TCBS plates were extracted separately. Qualitative PCR assays were performed on five colonies of each color per sampling site to confirm the presence of *Vibrio spp*.

For DNA extraction, colonies of the same color (yellow or green) were harvested from a plate and resuspended in 100 μL of nuclease-free water in a 2 mL microcentrifuge tube. The tubes were vortexed for 5 seconds and then heated to 95 °C in a heat block for 5 minutes. After heating, the tubes were centrifuged for 2 minutes at 12,000 rpm. The supernatant was placed in a labelled 1.5 mL microcentrifuge tube and stored at -80 °C until PCR.

Positive controls for PCR were generated from the American Type Culture Collection (ATCC; Manassas, VA) stock bacteria. For the *V. cholerae* (ATCC #14033) and *V. parahaemolyticus* (ATCC # 43996) cultures, 1 mL glycerol stock was placed in 9 mL Difco™ Nutrient Broth (VWR, Radnor, PA) in a loosely capped 15 mL tube and incubated at 37 °C for 24 hours. For the *V. vulnificus* (ATCC #27562) cultures, 1 mL glycerol stock was placed in 9 mL Difco™ Marine Broth (VWR, Radnor, PA) in a loosely capped 15 mL tube and incubated at 37 °C for 24 hours. After incubation, 1 mL of broth was pelleted down via centrifugation for 1 minute at 13,000 rpm. The supernatant was removed, and the pellet was resuspended in 100 mL nuclease-free water. The heat extraction method described above was used, and the DNA was stored at -80 °C until PCR.

PCR primer sets targeting the toxin gene of three *Vibrio* spp. of concern (Table S1) were used. A 25 µL reaction was assembled using 12.5 µL AccuStart^TM^ II PCR Supermix (Quantabio, Beverly, MA), 4 µL of forward primer (10 µM), 4 µL of reverse primer (10 µM), 2.5 mL of nuclease-free water, and 2 µL DNA template. For the non-template control, 2 µL of nuclease-free water was used instead of extracted DNA. Thermocycling conditions can be found in Table S2.

Thermocycling was completed on a MyCycler Thermal Cycler (Bio-Rad Laboratories, Hercules, CA). PCR products were visualized on a 1.5% agarose gel with a SYBR Safe DNA Gel Stain (Thermo Fisher Scientific, Waltham, MA). 10 µL of PCR product was loaded with 2 µL 6x Purple Gel Loading Dye (New England BioLabs, Ipswich, MA). A 100 bp DNA ladder (Promega, Madison, WI) was loaded in the same manner and used as a standard to estimate amplicon sizes.

### 2.5 Enumeration of *Vibrio* spp. using quantitative PCR (qPCR)

Undiluted water sample (100 mL) was filtered through 0.4 μm pore size, 0.47 mm diameter polycarbonate filters (MilliporeSigma, Burlington, MA) for use in qPCR. DNA extraction was performed using the DNeasy PowerLyzer PowerSoil kit (QIAGEN, Hilden, Germany) according to the manufacturer’s instructions. Briefly, bacterial cells collected on the filters were resuspended and homogenized in the Mini-Beadbeater-16 (Biospec Products, Bartlesville, OK) for 45 seconds. Following this, the sample underwent protein precipitation, inhibitor removal, DNA binding, and wash steps. DNA was eluted in 100 μL of DNase, RNase-free water and stored at -80°C until the qPCR assay was performed.

A SYBR Green assay with the 16S rRNA gene (Table S1) was used to determine concentrations of *Vibrio* spp. in water samples. Each 25 µL reaction consisted of 10 µL of 2 x Applied Biosystems™ PowerUp™ SYBR™ Green Master Mix (Thermo Fisher Scientific, Waltham, MA), 1 μL of 10 µM forward primer, 1 μL of 30 µM reverse primer, 4 μL of nuclease-free water, and 4 μL of sample or standard DNA. Thermocycling conditions are in Table S2.

Thermocycling and data collection was completed on the QuantStudio™ 3 Real-Time PCR System (Thermo Fisher Scientific, Waltham, MA). To develop standard curves for qPCR, A gBlock gene fragment (Integrated DNA Technologies, Coralville, IA) was designed to cover the 16S rRNA region of *Photobacterium angustum*. The gBlock was diluted 10-fold using nuclease-free water with concentrations ranging from 10^7^ to 10 gene copies/5µL. Additionally, each plate included non-template controls. All samples and standards were run in triplicate on a 0.2 mL, 96-well plate. The efficiency percentage and r^2^ values were computed for each standard curve with acceptable values between 90-110% for efficiency and ≥0.980 for r^2^. In this study, qPCR efficiency ranged from 93.3 to 95.9%, and r^2^ ranged from 0.996 to 1. Since *Vibrio* spp. contain between 9-14 copies of the 16S rRNA gene per cell, the number of presumed cells in each sample were calculated by dividing the gene copy number by 9.1. This number, 9.1, is the average number of 16S rRNA gene copies across *Vibrio* species (Johnson et al., 2010).

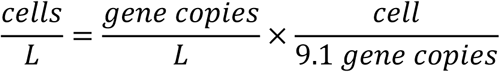

qPCR was also used to quantify the *Vibrio* spp. of concern (*V. cholerae*, *V. vulnificus*, and *V. parahaemolyticus*) using primers, probes, and gblocks as listed in Table S1. For specific *Vibrio* spp. qPCR assays, 20 μL reactions consisted of 10 μL 2x PerfeCTa qPCR Toughmix Low ROX (Quantabio, Beverly, MA), 1 μL of 10 µM forward primer, 1 μL of 10 µM reverse primer, and 0.2 μL of 10 µM probe, 3.8 μL of nuclease-free water and 4 μL of extracted DNA sample or standard DNA. Thermocycling conditions differed for each of the three species (Table S2). Thermocycling and data collection was completed on the QuantStudio™ 3 Real-Time PCR System (Thermo Fisher Scientific, Waltham, MA). All samples and standards were run in triplicate on a 0.2 mL, 96-well plate.

### 2.6 Genomic sequencing for culturable *Vibrio* spp. and bioinformatics

To determine the diversity of *Vibrio* spp. isolated from the estuary, bacterial colonies were harvested from TCBS Agar plates by using 1 mL of Luria-Bertani (LB) Broth (Thermo Fisher Scientific, Waltham, MA) and a sterile L-shaped loop to dislodge colonies from filter. Broth with suspended colonies was pipetted into a 1.5 mL cryotube and 150 μL autoclaved glycerol was added to make a 15% glycerol stock. Tubes were vortexed and stored at -80 °C. This process was completed in duplicate for each sample site.

For DNA extraction, the preserved cells from each site were pooled in 2-mL tubes and spun down for 5 minutes at 5,000 rpm. The supernatant was discarded, and the pellet was resuspended in ZymoBIOMICS DNA/RNA Shield solution (Zymo Research, Orange, CA). DNA was extracted according to the ZymoBIOMIC DNA/RNA Miniprep Kit DNA Purification protocol. In this study, 75 uL of the ZymoBIOMICS Microbial Community Standard was also extracted using the same protocol as a positive control.

DNA samples were prepared for sequencing with the Native Barcoding Kit NBD114.24 (Oxford Nanopore Technologies, Oxford, UK). For each sequencing, non-template controls and the ZymoBIOMICS Microbial Community Standard (Zymo Research, Orange, CA) as a positive control were also included. Tapestation electrophoresis (Agilent Technologies, Santa Clara, CA) was used to get the average input fragment length (kb) in the DNA library. This fragment length was used to calculate the volume of library that contained 50 fmol of DNA to load into a PromethION R10.4.1 Flow Cell (Oxford Nanopore Technologies, Oxford, UK) for nanopore sequencing with the MinKNOW v24.11.10 using the ‘fast’ Dorado Basecaller Setting. Samples were sequenced for up to 72 hours, the minimum quality score was set to 8 and minimum read length set to 1 kb.

Nanopore-generated FASTQ files were concatenated via the command line and uploaded to the IDseq mNGS Nanopore Metagenomic Pipeline v0.7. The pipeline filtered low-quality reads using fastp, removed host and human sequences with minimap2, subsampled to 1 million reads, assembled contigs using metaFlye, and aligned both contigs and unassembled reads to the NCBI nucleotide and protein databases using Minimap2 and DIAMOND (Buchfink et al., 2015; Chen et al., 2022; Kolmogorov et al., 2020; Li, 2018). Output reports included identified taxa, total bases per taxon, percent identity, alignment length, contig counts per taxon, and average E-value. Positive control samples were reviewed to confirm detection of organisms present in the microbial community standard. The results from the IDseq pipeline were uploaded to R (Version 2024.120 +467) for data visualization. Sequencing reads that aligned with *V. cholerae* genome in the IDseq were also uploaded to Cholerae Finder 1.0 (https://cge.food.dtu.dk/services/CholeraeFinder/) for serotyping. Reported species were screened to have a percent identification greater than 85%.

### 2.7 Statistical analysis

All statistical analyses were performed using R (Version 2024.120 +467). Data was cleaned using the dplyr and lubridate packages. The graphs were generated using ggplot2. A multiple linear regression model was used to assess the relationship between environmental factors (salinity, temperature, precipitation, dissolved oxygen, total dissolved solids, and pH) and *Vibrio* spp. concentrations collected via bacterial culture (CFU/L) and qPCR (cells/L). Values for bacterial culture concentration, qPCR concentration, and pH were log-transformed for normality. The equation for the linear regressions were as follows: log (Vibrio concentration) = β0+β1 (Salinity)+β2(Temperature)+β3(Precipitation)+β4(Dissolved Oxygen) +β5(log(pH))+ β6(total dissolved solids)+ɛ. The package “lmtest” was used to make linear models which were assessed for heteroscedasticity of residuals using the Breusch-Pagan test. Significance of environmental parameters on *Vibrio* spp. concentration was assessed using a type III ANOVA test where a p-value less than 0.05 was determined to be significant.

## 3 Results

### 3.1 Water quality and environmental parameters

Table 1 summarizes key water quality parameters of the estuary measured during the study. Water temperature ranged from 8.27 °C to 32.37 °C, with seasonal variation. Median water temperature during summer months (June, July, and August) was 30.5 °C, whereas during winter months (December, January, and February), median water temperature was 13.35 °C (Figure S1). Salinity ranged from 0.08 PSU to 10.01 PSU, which indicated brackish water. The dissolved oxygen percent saturation of water samples ranged from 51.7% to 111%. These values are based on expected solubility for a given salinity and temperature, but they can exceed 100% because of photosynthetic organisms that produce oxygen that do not immediately diffuse to reach equilibrium with the air. The highest precipitation value for the seven-day period before sample collection was 8.47 inches from Hurricane Francine in September 2024. Total dissolved solids ranged from 18 to 8484 mg/L throughout the course of the study. Of the samples that were tested, *E. coli* was detected in all but one sample, and enterococci was detected in all samples.

**Table 1.** Summary of key environmental parameters of Lake Pontchartrain collected over the course of the study.

| Parameter | Minimum | Maximum | Median |
| --- | --- | --- | --- |
| Temperature (°C) | 8.27 | 32.37 | 23.14 |
| Salinity (PSU) | 0.08 | 10.01 | 2.21 |
| Dissolved Oxygen (% saturation) | 51.70 | 111.00 | 87.90 |
| 7 Day Precipitation (in) | 0.00 | 8.47 | 0.50 |
| Total Dissolved Solids (mg/L) | 18 | 8484 | 1961.5 |
| *E. coli* (MPN/100 mL) | <1 | 1732.90 | 24.30 |
| *Enterococci* (MPN/100 mL) | 2 | >2419.6 | 79.80 |

### 3.2 Enumeration of *Vibrio* spp. in an estuarine lake

Using culture technique, *Vibrio* spp. was detected in 98% (n=99) of water samples, with bacterial counts ranged from 30 to 8.65 × 10^4^ CFU/L. Temporal variation in the prevalence of *Vibrio* spp. was observed over the course of the year (Figure 2). Concentrations of *Vibrio* spp. were greater during the summer months (June, July, and August, n=27), with the geometric mean of the concentrations being 3.2 × 10^4^ CFU/L. In the winter months (January and February, n=18), the average *Vibrio* concentration was 3.1 × 10^1^ CFU/L. A two-tailed t-test showed a significant difference between these seasons (p = 3.7 x 10^-6^). Quantitative PCR results showed a similar temporal pattern, with *Vibrio* DNA concentrations tending to be higher in water samples during the summer months (Figure 3). However, an exception was found in November in which water samples had the second highest geometric mean of *Vibrio* DNA (>3.0 × 10^4^ cells/L).

**Figure 2.**
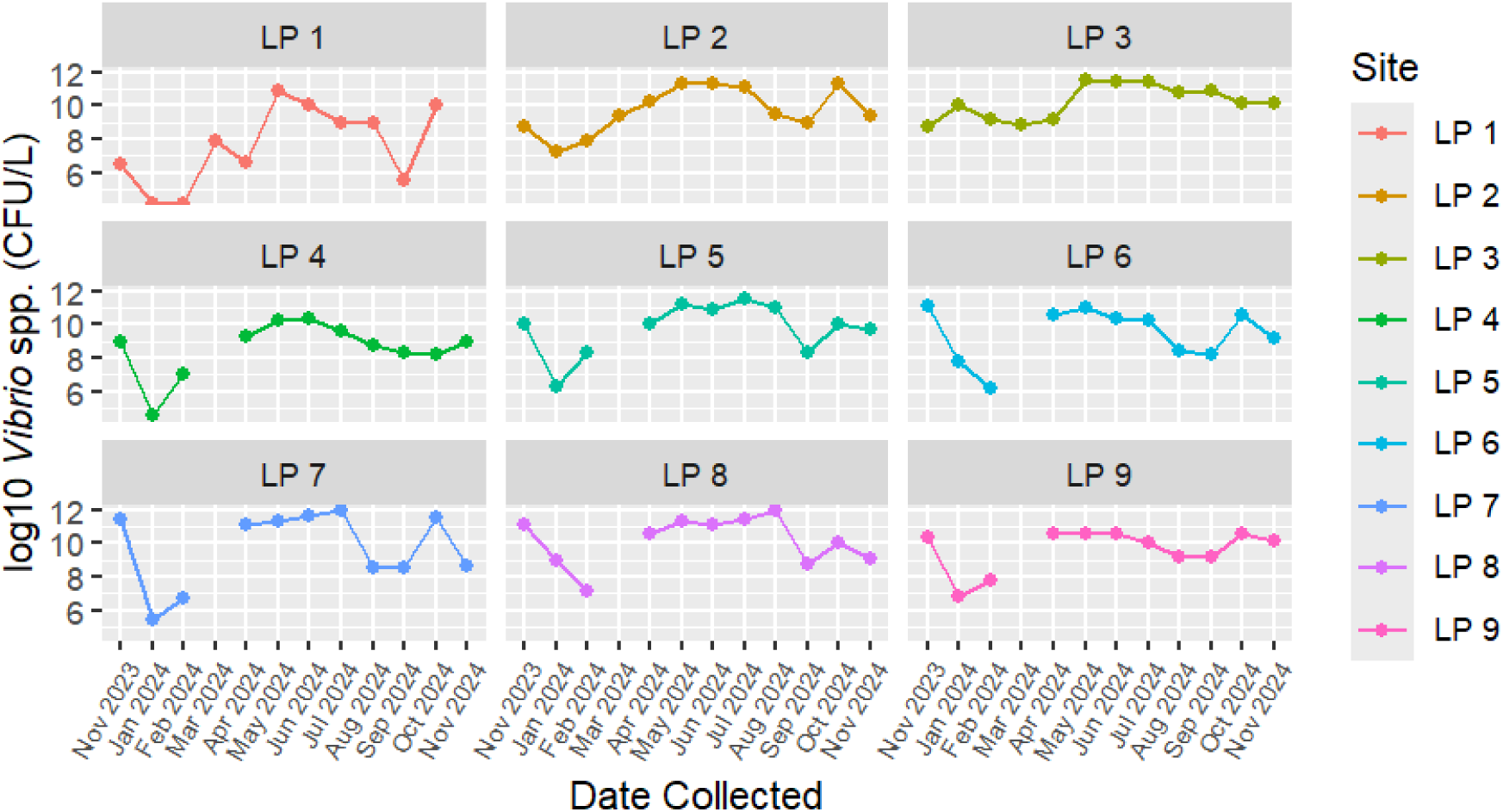
Temporal occurrence of *Vibrio* spp. in nine recreational sites as determined using culture assay with selective media.

**Figure 3.**
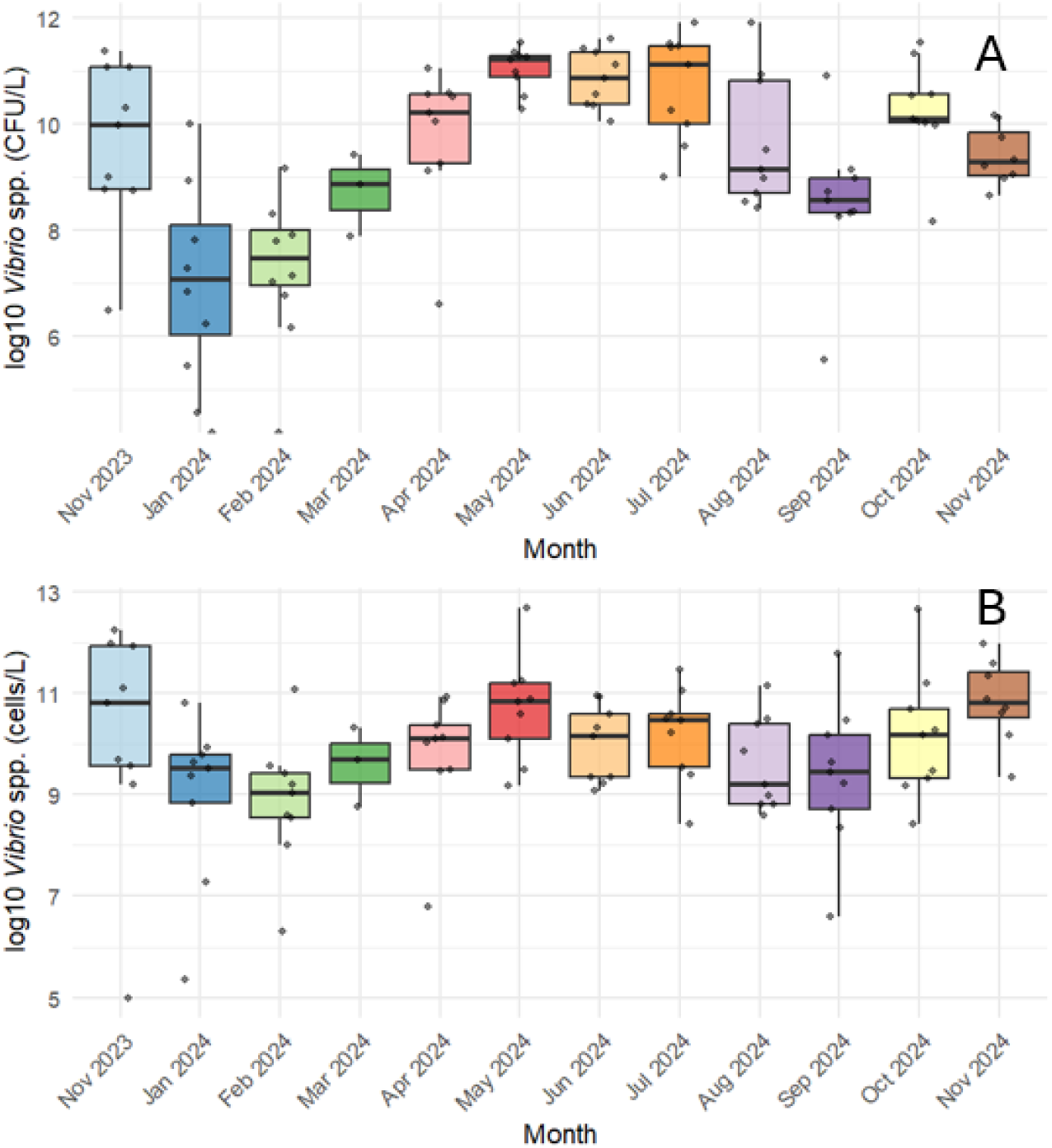
Monthly distributions and a comparison of (A) bacterial culture techniques and (B) qPCR quantification for the direct enumeration of *Vibrio* spp. in an estuarine lake.

The prevalence of *Vibrio* spp. also varied spatially in Lake Pontchartrain (Figure 4). For example, the sampling sites LP 1 and LP 4 had consistently lower geometric means of *Vibrio* spp., while sites LP 3 and LP 8 both had geometric means greater than 2.0 × 10^4^ CFU/L (Figure 4). The site LP 3 had the highest concentration of *Vibrio* DNA, which matched the trend for culturable *Vibrio*.

**Figure 4.**
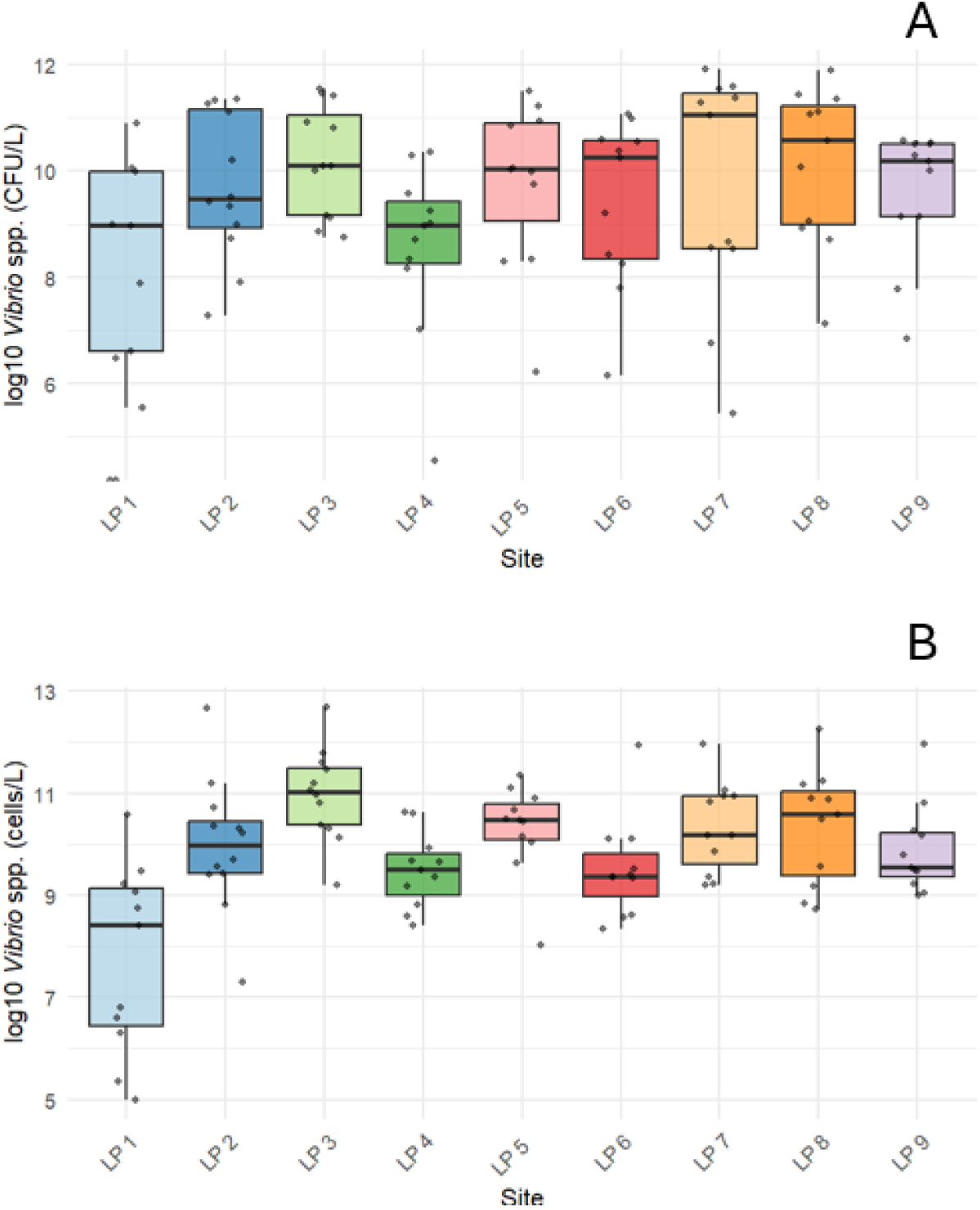
Spatial distributions and a comparison of (A) bacterial culture techniques and (B) qPCR quantification for the direct enumeration of *Vibrio* spp. in an estuarine lake over 12 months.

### 3.3 Associations between the occurrence of *Vibrio* spp. and water quality

Fecal indicator bacteria (FIB), such as, *E. coli* and enterococci, are commonly used as an indicator for fecal contamination in recreational water. In this study, the Pearson correlation test found very weak correlation (r < 0.2) between FIB and the detection of *Vibrios* in water samples. The months when average FIB concentrations were at their peak were not the same months when *Vibrio* spp. concentrations were highest (Table S3 and Figure 3).

The linear models between *Vibrio* concentrations with environmental factors showed that salinity and temperature were significant predictors of *Vibrio* concentrations from both bacterial culture and direct qPCR methods (Table 2). Additionally, precipitation was a significant predictor for culturable *Vibrio* spp. in this study. The model for bacterial culture concentration had a higher adjusted r^2^ value compared to the qPCR model (0.3914 and 0.1401, respectively), meaning that concentrations from bacterial culture methods were better predicted by environmental factors.

**Table 2.**
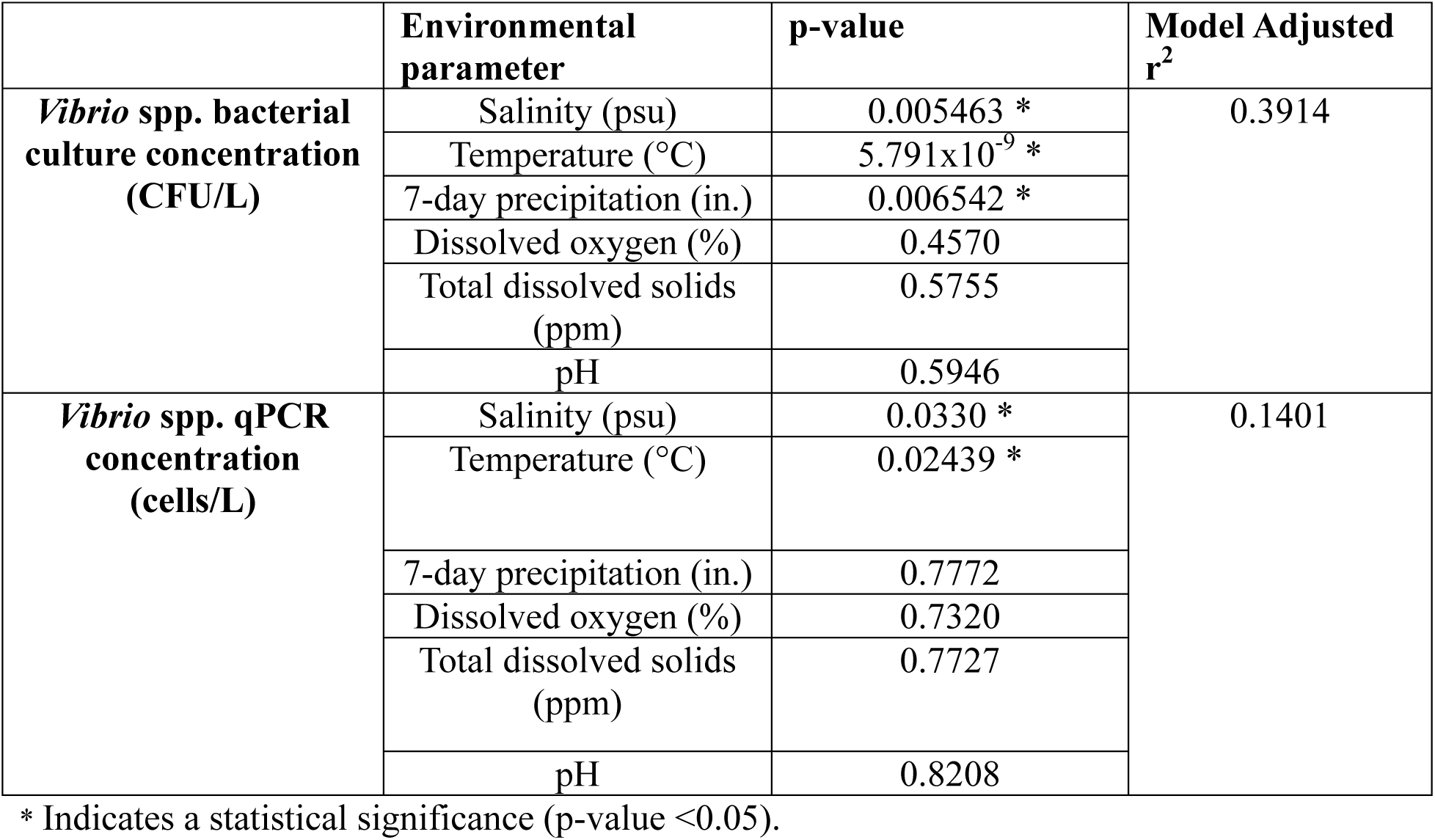
Linear models comparing *Vibrio* quantification and environmental parameters.

Furthermore, a Pearson correlation test shows that *Vibrio* concentrations as determined using culture method and direct qPCR were weakly correlated with an r^2^ value of 0.2585.

Additional temporal analysis focused on samples collected from the south shore sites (LP4 – LP9) in September 2024, the week following Hurricane Francine, revealed that several environmental parameters were significantly different compared to samples collected at the same sites later in Fall 2024. Specifically, salinity, rainfall, and temperature differed significantly between September and the months of October and November 2024 (p < 0.05). However*, Vibrio* spp. concentrations, as measured by both bacterial culture and qPCR methods, were not significantly different across these time points (p > 0.05). In contrast, a parametric t-test showed a significant difference in culturable *Vibrio* concentrations when comparing water samples collected in November 2024 to those collected in November 2023, a period when no named storms impacted the New Orleans area. Average *Vibrio spp*. concentrations measured by culture were 5.1 × 10⁴ CFU/L in November 2023 and 1.2 × 10⁴ CFU/L in November 2024, a statistically significant difference (p = 0.047).

### 3.4 Presence of *Vibrio* species of concern

Conventional PCR targeting the *toxR* gene, a transcriptional regulator of virulence genes, was used to confirm and identify culturable *Vibrio* species of concern. Among the samples collected (n = 101), 85.2% (n = 86) tested positive for *V. cholerae,* 20.8% (n=21) for *V. parahaemolyticus*, and 50.5% (n=51) for *V. vulnificus toxR* genes (Figure 5, Table S4). Notably, nine samples (8.9%) were positive for all three species. In June, July, September, and October, all sampling sites tested positive for the *V. cholerae toxR* gene (Figure 5).

**Figure 5.**
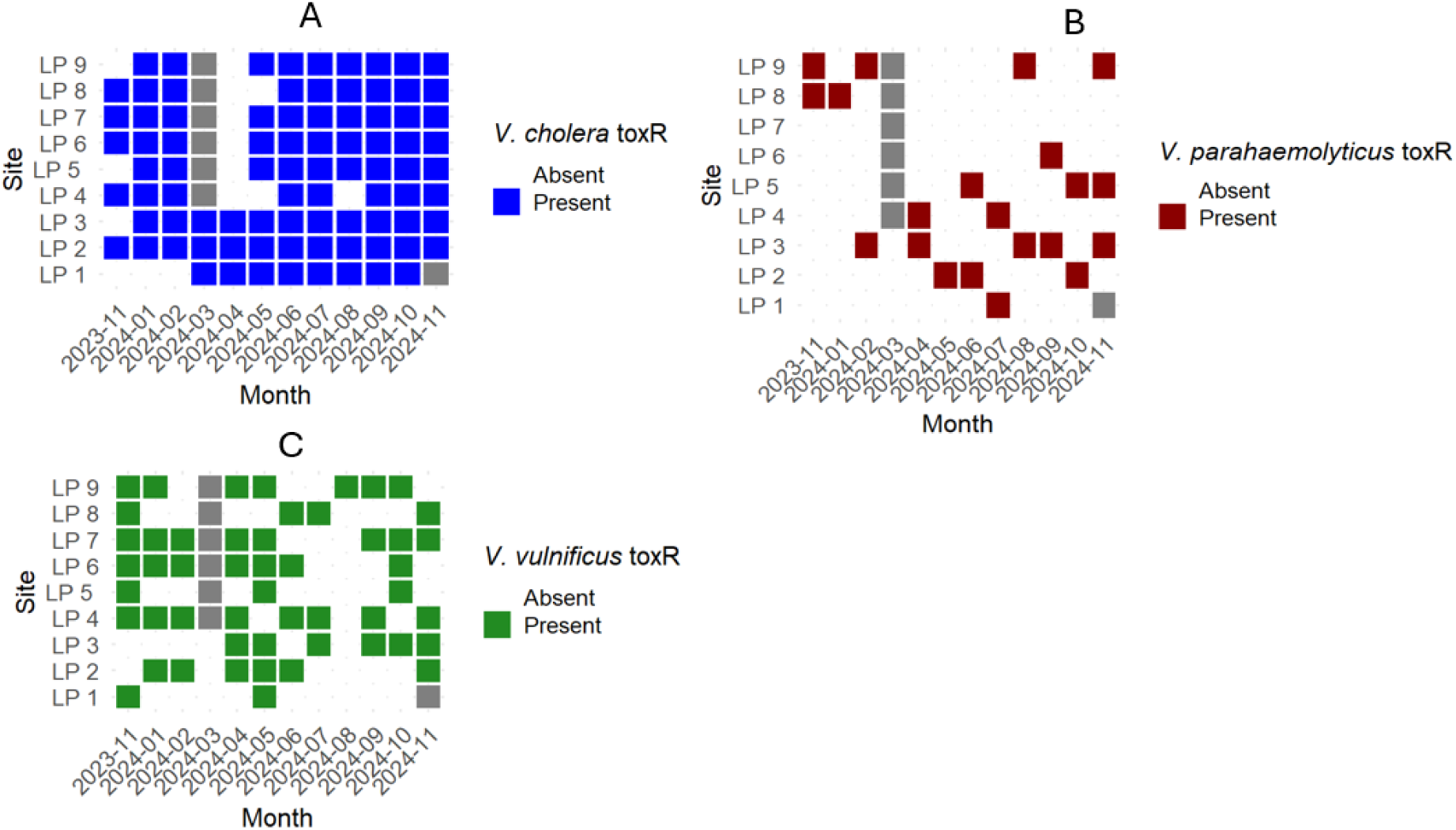
Spatiotemporal variability of culturable (A) *V. cholerae*, (B) *V. parahaemolyticus*, and (C) *V.* vulnificus in an estuary as confirmed using conventional PCR targeting the *toxR* gene. Grey squares represent time periods when samples were not collected.

Quantitative PCR assays were also used for the direct detection of *Vibrio* species of concern (Table S4). In contrast to the detection of *toxR* gene, the *V. cholerae ctxA* gene, which codes for the A subunit of the cholera toxin, was only found in four samples (3.9%), with concentrations ranging from 33.24 to 105.8 gene copies per liter (GC/L) (Figure 6 and Table S4). The *V. parahaemolyticus gyrB* gene, which codes for the DNA gyrase specific to this species, was detected in 41.6% of samples, with concentrations ranging from 12.28 to 1.28 x 10^4^ GC/L. With direct qPCR, there was no clear trend in times of year or specific sites where *V. parahaemolyticus gyrB* gene were more prevalent (Figure 6 and Table S4).

**Figure 6.**
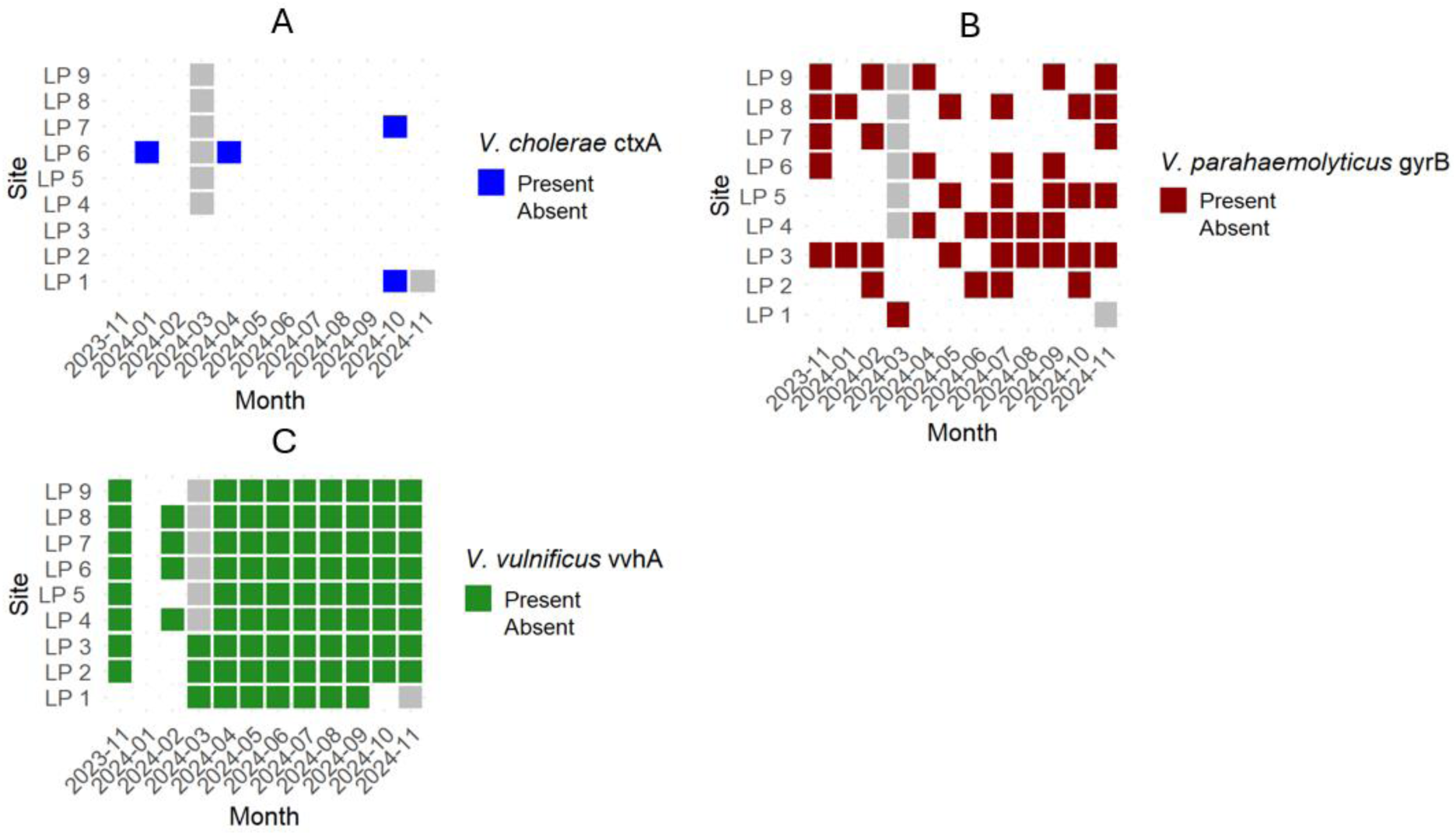
Spatiotemporal variability of (A) *V. cholerae ctxA* gene, (B) *V. parahaemolyticus gyrB* gene, and (C) *V. vulnificus vvhA* gene in an estuary as confirmed by qPCR. Grey squares represent time periods when samples were not collected.

Direct qPCR revealed a high prevalence of the *V. vulnificus vvhA* gene, which encodes a hemolysin protein associated with pathogenicity. The gene was detected in 84.2% (n = 85) of samples, with concentrations ranging from 6.49 to 7.21 × 10⁵ GC/L (Figure 6, Table S4).

Temporal patterns were evident: no *vvhA* gene was detected in January 2024, while all samples from April through September 2024 were tested positive (Figure 6). The highest concentrations occurred from June to September, ranging from 9.23 × 10^2^ to 7.21 × 10⁵ GC/L (Figure 7A, Table S5). Spatially, the highest concentrations were observed at sites LP 4, LP 6, LP 7, and LP 8—all located on the south shore of Lake Pontchartrain (Figure 7B, Table S6).

**Figure 7.**
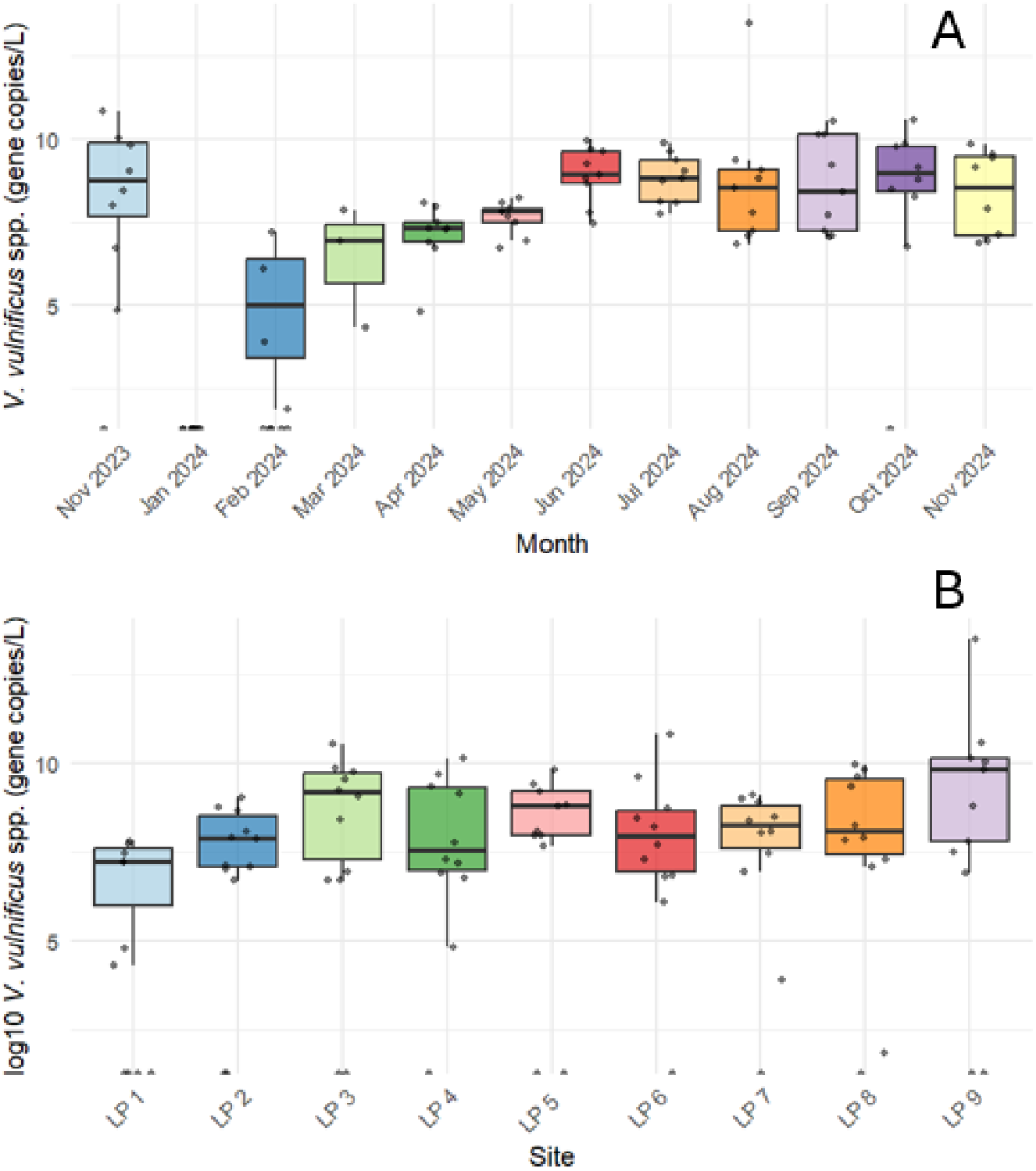
The concentration of *V. vulnificus* hemolysin (*vvhA*) gene in an estuary. Quantitative PCR showing (A) temporal and (B) spatial distributions of the gene.

### 3.5 Diversity of *Vibrio* spp. in an estuarine lake

Four sequencing libraries were created from the 60 different samples that were barcoded for a long-read nanopore sequencing. For sequencing runs that failed before 72 hours, the same library was run again to generate more reads. In total, 7 runs were used in this study, with a total of 146 million reads generated and 534.86 Gb of DNA sequenced (Table S7). The N50 values for the reads ranged from 4.89 to 8.31 kb.

Sequencing bacterial colonies and comparing them to known sequences in the NCBI GenBank database revealed that the majority of the reads aligned with *V. cholerae* genomes (Figure 8). Of the total sequencing reads obtained, 56.4% were identified as *V. cholerae*, followed by *V. mimicus* with 21.5%, and *V. vulnificus* with 11.9%. Other *Vibrio* spp. identified by sequencing included *V. metoecus* and *V. parahaemolyticus*. The “Other” category contains those species contributed to less than 1% of the sequenced nucleotides, which includes over 100 other known *Vibrio* species (Figure 8 and Table S8). Of the reads that aligned with *V. cholerae* genomes (n=60), 14.8% of these samples contained the *wbf* gene cluster indicative of the O139 serotype (Figure 8).

**Figure 8.**
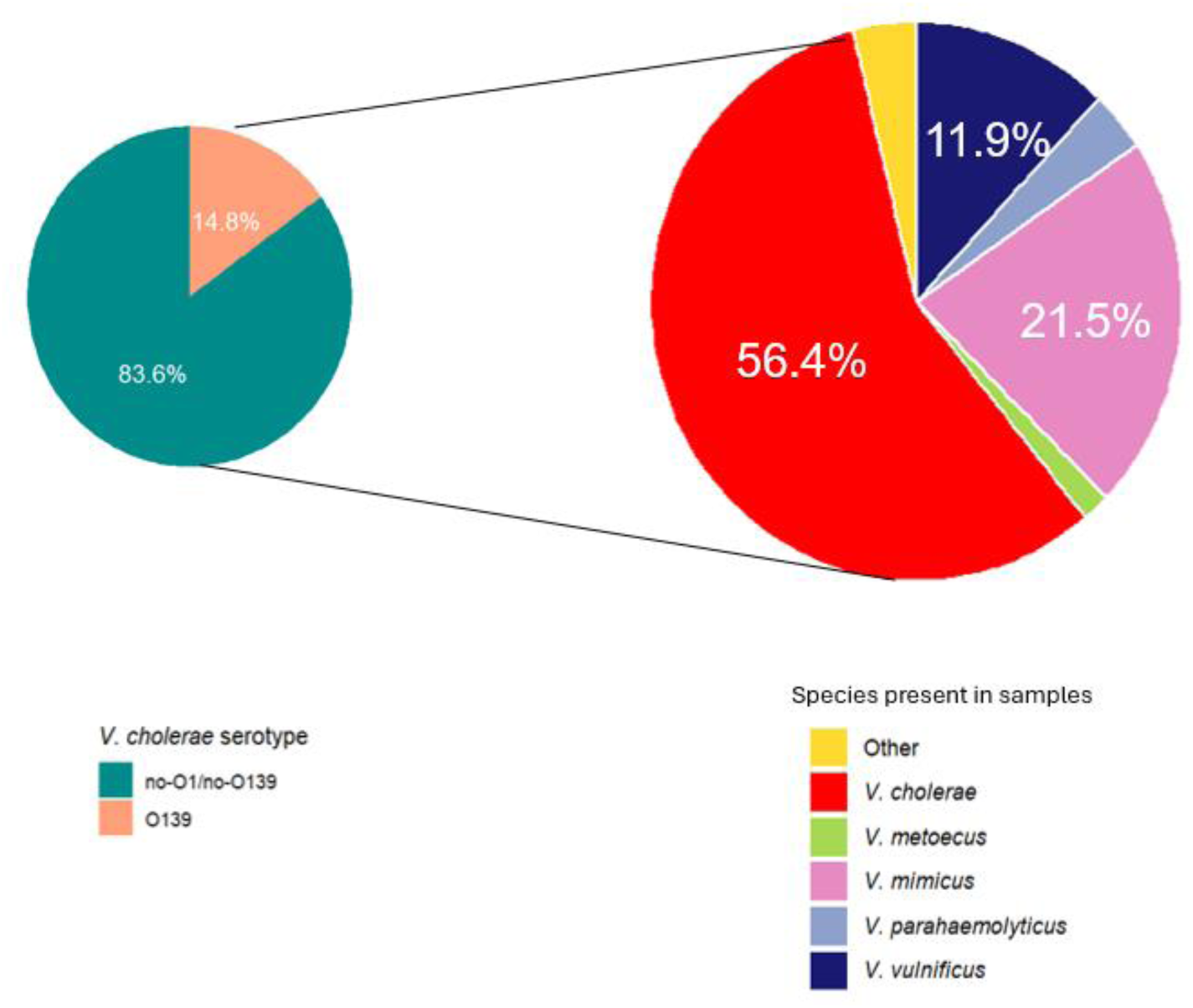
Proportions sequenced nucleotides that correspond to each *Vibrio* species. *Vibrio cholerae* made up the majority, with 14.8% of the samples that were identified as *V. cholerae* containing genes from the O139 serotype.

## 4 Discussion

Previous studies in Louisiana have focused on *Vibrio* spp. infections (Thomas et al., 2007) or culturable *Vibrio* spp. in oysters from the Gulf of Mexico (Han et al., 2007). This study instead focused on the occurrence and diversity of *Vibrio* spp. in brackish water from Lake Pontchartrain, specifically in recreational areas. At these sites, mean culturable *Vibrio* spp. concentrations were found to be highest during the months of May, June, and July, which aligns with results from the linear model comparing these concentrations and environmental factors.

The linear model showed that temperature, salinity, and precipitation were significant predictors of culturable *Vibrio* concentration. This is in line with other studies of environmental *Vibrio* that have noted a linear relationship between temperature and *Vibrio* concentrations (Brumfield, Chen, et al., 2023; Deeb et al., 2018; Green et al., 2019).

Similarly, a linear model showed that both temperature and salinity were significant predictors of *Vibrio* 16S rRNA gene concentrations. However, the r^2^ values for these models show that all environmental factors included in the model explained about 39% of the culturable *Vibrio* spp. variation and only about 14% of the qPCR *Vibrio* concentration. It is possible that gene concentrations would be less influenced by environmental factors as *Vibrio* spp. can exist in a dormant, nonculturable state when environmental conditions fluctuate (Colwell & Huq, 1994). *Vibrio* spp. can survive at temperatures as low as 4 °C, but optimum growth occurs at temperatures between 25 °C and 30 °C. It is possible that this VBNC state contributed to the lower r^2^ value for the qPCR model, as this method would account for bacteria in a metabolically dormant state during the winter months (Sheikh et al., 2022). Environmental factors such as salinity, temperature, and precipitation significantly influenced *Vibrio* communities in this estuarine system. However, because less than half of the variation in culturable and qPCR-detected *Vibrio* was explained by these factors, they alone are insufficient for predicting health risks to recreators. These results show the importance of monitoring for *Vibrio* spp. in recreational waters throughout the year. Even with all six environmental factors included in the model (salinity, temperature, precipitation, dissolved oxygen, total dissolved solids, and pH), less than half of the fluctuation in culturable *Vibrio* spp. concentrations were explained.

Many recreational sites were tested for FIB and held to the EPA recommendation of less than 35 *Enterococci* per 100 mL and less than 126 *E. coli* per 100 mL of water sample. Since cholera can also be spread through fecal matter, FIB concentrations were also collected for comparison (EPA, 2012). Fecal indicator bacteria concentration in many of the samples collected in this study exceeded these levels, but there was a weak correlation between *Vibrio* spp. concentration and FIB. This again underscores the need for *Vibrio* testing, as environmental factors or FIB presence cannot fully predict the variability of these bacteria.

In addition to normal seasonal variations in environmental parameters, Hurricane Francine impacted the city of New Orleans during September 2024. This event caused significant differences between salinity, rainfall, and temperature for this month compared to other samples collected during the Fall of 2024. A comparison between November 2024, after this extreme weather event, and the previous November when New Orleans experienced no named tropical disturbances, shows that the *Vibrio* spp. was significantly lower in November 2024. This differs from a study conducted after Hurricane Ian made landfall in Florida in 2022, which showed that environmental changes due to the storm made the waters more favorable for *Vibrio* spp. growth (Brumfield, Usmani, et al., 2023). However, it is not possible to compare *Vibrio* spp. concentrations from September 2023 to determine if *Vibrio* spp. concentrations may have been higher immediately following Hurricane Francine. While in this study, comparison could only be made two months after the hurricane, it is important to continue to study how these extreme weather events can influence *Vibrio* spp. populations.

For the samples collected in this study, *V. cholerae toxR* gene was present in most samples, followed by *V. vulnificus toxR* gene, and finally *V. parahaemolyticus toxR* gene. This is corroborated by past research on water samples from Lake Pontchartrain and the Gulf of Mexico where these three species of *Vibrio* were present (Johnson et al., 2010; Nigro et al., 2011).

However, epidemiological data suggests that *V. parahaemolyticus* is the most commonly reported cause of vibriosis in Louisiana and the U.S. (CDC, 2019; Newton et al., 2012; *Vibrios Annual Report*, 2018), but *V. parahaemolyticus toxR* gene was only present in 14% of samples. This could be attributed to that *V. parahaemolyticus* tends to be more associated with shellfish, particularly oysters while *V. cholerae* is found more often in the water column (Jones et al., 2014). While *V. parahaemolyticus* was found in fewer water samples, its association with seafood could explain the higher rates of infection.

Quantitative PCR results for specific toxin genes of *V. cholerae* and *V. vulnificus*, cholera toxin and *V. vulnificus* hemolysin respectively, show that these are not directly correlated to presence of the *toxR* gene in either of these species. In this study, the cholera toxin gene (*ctxA*) was only found in 4% of samples. However, in 128 samples collected in Lake Pontchartrain between 2005 and 2006, none tested positive for the *ctxA* gene (Nigro et al., 2011). This could be due to changing conditions in the lake or because in this study the *ctxA* gene was identified using qPCR, which can identify the gene even if the bacteria are in a VBNC state.

While both the *V. vulnificus* hemolysin gene and the *Vibrio* 16S rRNA concentrations peaked during summer months, they did not peak at the same time. This could mean that risk of interaction with toxigenic *Vibrio* spp. persists in months when total *Vibrio* populations are lower.

This means that even in times when there are lower concentrations of *Vibrio* bacteria in the water, recreators still interface with toxin-producing species. Additionally, the site with the highest concentration of both culturable *Vibrio* spp. and *Vibrio* 16S rRNA genes, LP 3, did not contain the highest concentration of *V. vulnificus* hemolysin genes. This shows that the spatial distribution of pathogenic *Vibrio* could differ from the overall distribution of *Vibrio* species.

Analysis of sequences revealed that the *V. cholerae* made up the largest proportion of culturable *Vibrio* spp. in collected samples and the O139 serotype was present at sampling sites. This strain emerged in the 1990s, and has been responsible for global epidemic outbreaks of *V. cholerae* (Faruque et al., 2003). Cholera infections make up a very small proportion of total reported *Vibrio* infections in the U.S. (CDC, 2019), and there has only be one confirmed *V*.

*cholerae* O139 case in Louisiana since 1988 (*Vibrios Annual Report*, 2018).The O139 strain does produce the cholera toxin that can cause watery stool and potentially deadly dehydration (Faruque et al., 2003). Furthermore, sequencing of the cultured *Vibrio* colonies revealed 100 of species of *Vibrio* including *V. mimicus*, which is very similar to *V. cholerae* and can contain the genes for the cholera toxin (Wang et al., 2011; Zhou et al., 2021). While much of the work on *Vibrio* spp. focuses on the three species of concern, *V. cholerae, V. vulnificus, and V. parahaemolyticus*, there is a large diversity of bacteria in this genus, and it could be important to look at the dynamics between different species beyond those of concern to humans.

One advantage of bacterial isolate genome sequencing is that it does not require primer design like target-specific PCR. This allows for the detection of all *Vibrio* spp. and serotypes in a sample, not just the pre-defined species of concern (Chen et al., 2022). Furthermore, the use of nanopore long-read sequencing allows for the identification of specific virulence genes, like the *wbz* gene specific to the *V.* cholerae O139 serotype, that can help in understanding the *Vibrio* spp. community beyond the species level. However, this method is expensive and time consuming as compared to more rapid detection with PCR. Although culture methods have long been considered the gold standard for pathogen detection, they require incubation times between 24 to 48 hours, which can hinder rapid detection of *Vibrio* spp. in recreation waters. Since qPCR can be run without growing the bacteria, it can get faster results, but it tests for the presence of DNA rather than viable cells (Zampieri et al., 2021). This means that the qPCR technique can quantify all living *Vibrio* in a sample, even if they are metabolically dormant, but it also quantifies genetic material present in dead cells. These approaches all have their advantages and disadvantages, but they can be used together to get a comprehensive understanding of microbial communities in recreational water (Zampieri et al., 2021).

The presence of *Vibrio* spp. at recreational sites along the shore of Lake Pontchartrain is a public health concern for recreators who use the water for swimming, boating, and fishing. While culturable *Vibrio* spp. concentrations peak in the summer, the bacteria are present in recreational waters throughout the year, and variance cannot be fully explained by environmental factors.

Toxigenic *V. cholerae*, *V. vulnificus*, and *V. parahaemolyticus* as well as other *Vibrio* spp. can infect people who accidentally ingest the water or expose an open wound and severe cases can lead to hospitalization or even death (CDC, 2019).

## 5 Conclusion

*Vibrio* spp. concentrations were found to be highest in recreational water of Lake Pontchartrain in the summer, with a moderate correlation between water temperature and culturable *Vibrio* spp., but the bacteria persisted throughout the year. Furthermore, PCR and genome sequencing revealed the presence of *V. cholerae* including the O139 serotype, *V. vulnificus*, *V. parahaemolyticus*, *V. mimicus* and hundreds of other species in these samples. It would be valuable for future studies to determine other environmental parameters that influence the variation in *Vibrio* spp. populations and the interactions between the different bacterial species that make up the *Vibrio* community in these waters. This study has broader public health and environmental implications as future climate scenarios suggest that summers will become longer and warmer, increasing exposure risk of pathogenic *Vibrio* during recreational activities (Deeb et al., 2018; Froelich & Daines, 2020; Leal Filho et al., 2022).

## Acknowledgements

Funding for this study was provided by grants from the Louisiana Sea Grant Undergraduate Research Opportunities Program and Tulane University Center for Engaged Learning & Teaching. We thank Dr. Huiyun Wu and Keegan Brighton for assisting in data analysis and experimental design. We also thank Dr. David Mullin and Dr. Tim McLean who gave their valuable insight into this research and manuscript.

## CRediT authorship contribution statement

**Annika Nelson:** Formal analysis, Investigation, Methodology, Validation, Visualization, Writing – original draft. **Fernanda Mac-Allister Cedraz:** Formal analysis, Supervision, Investigation, Methodology, Writing – review & editing. **Katie Vigil:** Investigation, Methodology, Supervision, Formal Analysis, Writing – review & editing. **Joshua Alarcon:** Investigation, Methodology, Writing – review & editing. **Tiong Gim Aw**: Conceptualization, Investigation, Funding acquisition, Methodology, Project administration, Resources, Supervision, Writing – review & editing.

## Data Availability

The raw data required to reproduce the above findings are available to be shared.

